# Sex differences in gene regulation in the dorsal root ganglion after nerve injury

**DOI:** 10.1101/152652

**Authors:** Kimberly E. Stephens, Weiqiang Zhou, Zhicheng Ji, Shaoqiu He, Hongkai Ji, Yun Guan, Sean D. Taverna

## Abstract

Pain is a subjective experience derived from complex interactions among biological, environmental, and psychosocial pathways. Sex differences in pain sensitivity and chronic pain prevalence are well established. However, the molecular causes underlying these sex dimorphisms are poorly understood particularly with regard to the role of the peripheral nervous system. Here we sought to identify shared and distinct gene networks functioning in the peripheral nervous systems that may contribute to sex differences of pain after nerve injury. We performed RNA-seq on dorsal root ganglia following chronic constriction injury of the sciatic nerve in male and female rats. Analysis from paired naive and injured tissues showed that 1456 genes were differentially expressed between sexes. Appreciating sex-related gene expression differences and similarities in neuropathic pain models may help to improve the translational relevance to clinical populations and efficacy of clinical trials of this major health issue.

## INTRODUCTION

Once established chronic pain is often resistant to existing treatments and associated with adverse health outcomes such as decreased quality of life (2009), alterations in mood (Schaefer et al. 2014) and sleep patterns (Cheatle et al. 2016), and disability (Global Burden of Disease Study 2015). Sex differences in the susceptibility to most chronic pain conditions are well established (Fillingim et al. 2009) and experimental pain studies consistently demonstrate greater pain sensitivity, increased pain ratings, and decreased tolerance to a variety of pain modalities in women versus men (Torrance et al. 2006; Bouhassira et al. 2008; Fillingim et al. 2009; Mogil 2012; Bartley and Fillingim 2013). However, the mechanisms underpinning these sex dimorphisms are poorly understood.

While the inclusion of women in clinical trials has increased dramatically during the past 20 years, a strong bias towards the exclusive use of male animals in preclinical studies of neuropathic pain has persisted (Mogil and Chanda 2005; Beery and Zucker 2011). The limited use of females in mechanistic studies of neuropathic pain has obfuscated our understanding, management, and treatment of neuropathic pain in either sex. The few studies that have used rodent models of chronic pain to examine sex-specific differences have primarily focused on the hormonal and neuroimmune effects on pain modulation in the central nervous system (CNS) (Rosen et al. 2017). For example, activated immune cells (e.g., microglia) in the spinal cord dorsal horn release inflammatory mediators (e.g., cytokines) in response to tissue damage which promote neuronal excitability (Grace et al. 2014). The role of this microglia-neuronal signaling pathways in pain pathophysiology had been well-characterized using predominantly male mice; however, recent evidence now demonstrates the use of distinct immune cell types to mediate pain behaviors in female versus male mice (Sorge et al. 2011; Sorge et al. 2015).

Chronic pain following peripheral nerve injury is associated with profound changes in gene expression that alter synaptic plasticity, neuroimmune interactions, and long term potentiation. Unfortunately, the majority of studies have examined gene expression changes exclusively in males (Mogil and Chanda 2005; Beery and Zucker 2011; LaCroix-Fralish et al. 2011). One notable exception is a 2006 study by LaCroix-Fralish and colleagues (LaCroix-Fralish et al. 2006) in which microarrays were used to measure the gene expression in the rat dorsal horn after spinal nerve ligation in male and female rats. Despite the potential for gene expression to yield insight into the molecular underpinnings of chronic pain, a transcriptome-wide assessment of gene expression in the peripheral nervous system (PNS) after nerve injury in male and female rats has not been reported.

Here we sought to identify the similarities and differences in gene regulation between male and female rats in the PNS following peripheral nerve injury. We compare large-scale transcriptome analyses of dorsal root ganglia (DRGs) from male and female rats, in both naive and a peripheral nerve injury model, and identified differential expression of mediators of neurochemical mechanisms. In particular, our RNA-seq data shows that genes which facilitate synaptic transmission in naïve and injured females were not expressed in males. Therefore, sexually dimorphic roles of sensory neurons and glia in the PNS may be an important consideration in clinical drug trials designed to evaluate treatments for chronic pain.

## RESULTS

RNA-seq transcriptome profiles of DRGs from gonadally intact, adult male and female rats were measured from naïve animals and animals at 14 days after sciatic chronic constriction injury (CCI) (Figure 1). Basic biological influences that regulate responses to nociceptive stimuli and the generation of chronic pain have largely been investigated using neuropathic pain models in rodents. In an effort to isolate the pain behaviors from other experimental procedures used to create the model the majority of these prior studies used sham procedures as a comparison group. In sham procedures, all experimental procedures are performed with the exception of nerve injury from mechanical manipulation or administration of an active compound. In the present study, we use naïve animals as a comparison group so that we may capture all changes that are associated with clinical neuropathic pain conditions (i.e., skin incision, damage to nerve terminals, deep muscle tissue with associated inflammatory response).

**Figure 1:**
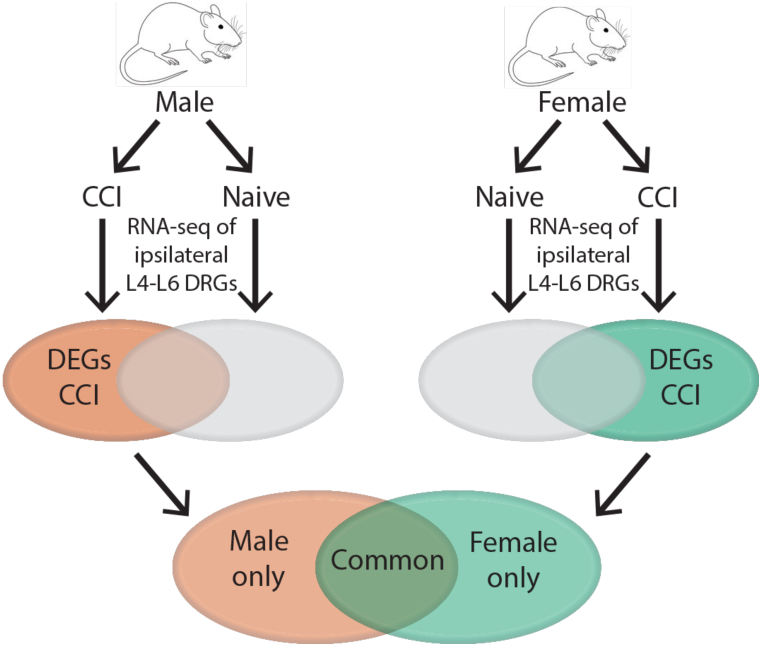
Schematic diagram of experimental procedures. Male and female rats were randomly assigned to the naïve group or receive CCI. Total RNA was isolated from the L4-L6 dorsal root ganglia (DRG) of naïve rats and on day 14 after chronic constriction injury (CCI) to the sciatic nerve. Libraries were constructed after poly(A) selection and sequenced. RNA-seq was performed on ipsilateral L4-L6 DRGs from each animal. Differentially expressed genes (DEG), defined as genes expressed after CCI versus naïve with a |log_2_fold change (FC)| > 0.5 and an adjusted p-value < 0.05 were compared between male and female rats.

### Differentially regulated genes in DRGs of naïve male and female rats

To identify sex-specific differences of gene expression in DRG under un-injured basal conditions, we performed RNA-seq and measured all poly(A)-containing transcripts expressed in the L4-L6 DRG of naïve rats (Figure 2A). Expression levels of 20,011 genes were evaluated and the majority (19,093 genes; 95.4%) showed no significant difference in expression level between males and females (Figure 2B). Of the 918 genes with expression levels that differed significantly between males and females, 719 (78.3%) showed increased in expression in males versus females and 199 (21.7%) showed increased expression in females versus males (Figure 2C-D). Lists of these genes with increased and decreased expression in the DRGs between male and female rats are provided as Supplemental Table 2 and Supplemental Table 3. These male and female gene lists were then subject to gene ontology analysis using Metascape. Gene ontology (GO) pathways associated with genes with increased expression in the male naive DRG include RNA splicing (GO:0008380), skeletal system morphogenesis (GO:0048705), and bone trabecular morphogenesis (GO:0061430) (Figure 2E). GO pathways associated with genes with increased expression in the female naive DRG were different from that in males and include antigen processing and presentation of peptide antigen via MHC class II (GO:0002495), inorganic ion transmission transport (GO:0098660), and negative regulation of endopeptidase activity (GO: 0010951).

**Figure 2:**
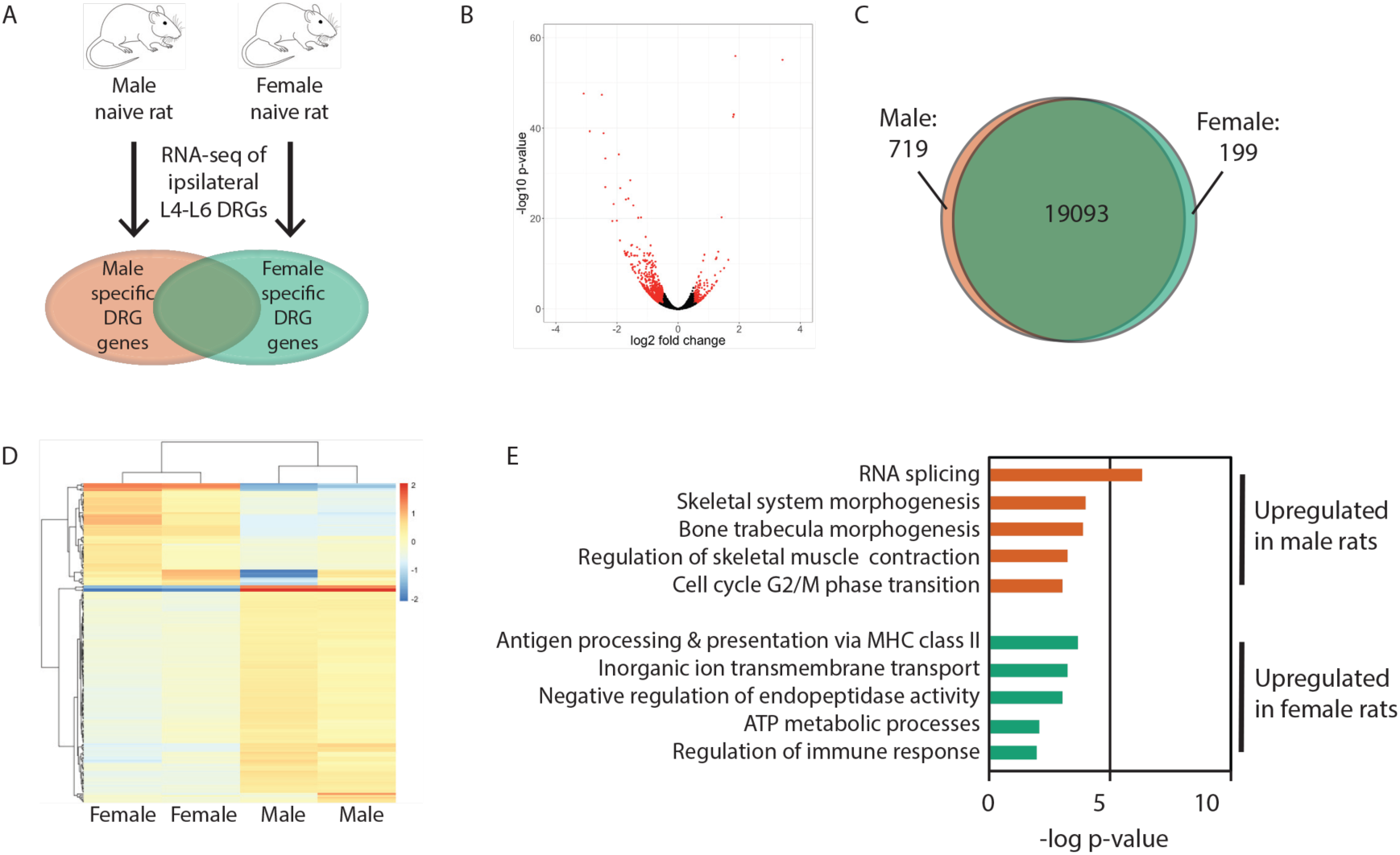
Sex differences of gene expression in the dorsal root ganglia (DRG) from naïve rats. A) Schematic diagram of experiment. RNA-seq performed on ipsilateral L4-L6 DRGs from naïve male and female rats. Differentially expressed genes (DEG) in one sex are identified by a |log_2_fold change (FC)| > 0.5 and an adjusted p-value < 0.05. B) Volcano plot of differential gene expression in naïve rats between male and female rats. Significant genes are designated in red. Fold change represents the ratio of gene expression in female to male rats. C) Venn diagram representing the number of genes expressed in males (red), females (green), and in both sexes (overlap). D) Heatmap that show the log_2_FC of the 200 most variable genes between naïve male and naive female rats. E) Gene ontology pathways associated with increased expression of genes in male (top) and female (bottom) naïve DRG.

### Differentially regulated genes in the dorsal root ganglion after nerve injury in male rats

To determine the effects of peripheral nerve injury on gene expression in the DRGs of male rats, we evaluated RNA-seq data obtained 14 days after CCI and compared these expression profiles to those from naive males (Figure 3A). Compared to naive DRGs, DRGs from male CCI rats differentially expressed 3316 genes (16.3%) (Figure 3B). Of these 3316 differential genes, 1429 genes (43.1%) were upregulated after injury and 1887 genes (56.9%) were downregulated after injury (Figure 3B). Despite the use of naïve rats as the comparison group, we found that a large number of upregulated genes after CCI (e.g., *Reg3b*, *Vgf*, *Atf*, *Cacna2d1*, *Gal*, *Npy*, *Gap43*) are the same as those previously reported to be upregulated in the DRG after nerve injury which used sham-operated rats as the comparison group (Costigan et al. 2010; LaCroix-Fralish et al. 2011). The genes with the largest log_2_fold increase and log_2_fold decrease in gene expression following CCI are listed in Table 1 and Table 2, respectively. GO analysis of the upregulated transcripts confirmed significant enrichment among nerve injury and pain related processes (Figure 3C). Among the 719 genes that showed increased expression in DRGs of naïve male rats compared to female naïve rats, 438 genes (61%) were downregulated following CCI whereas only 23 genes (3.2%) were upregulated following CCI.

**Figure 3:**
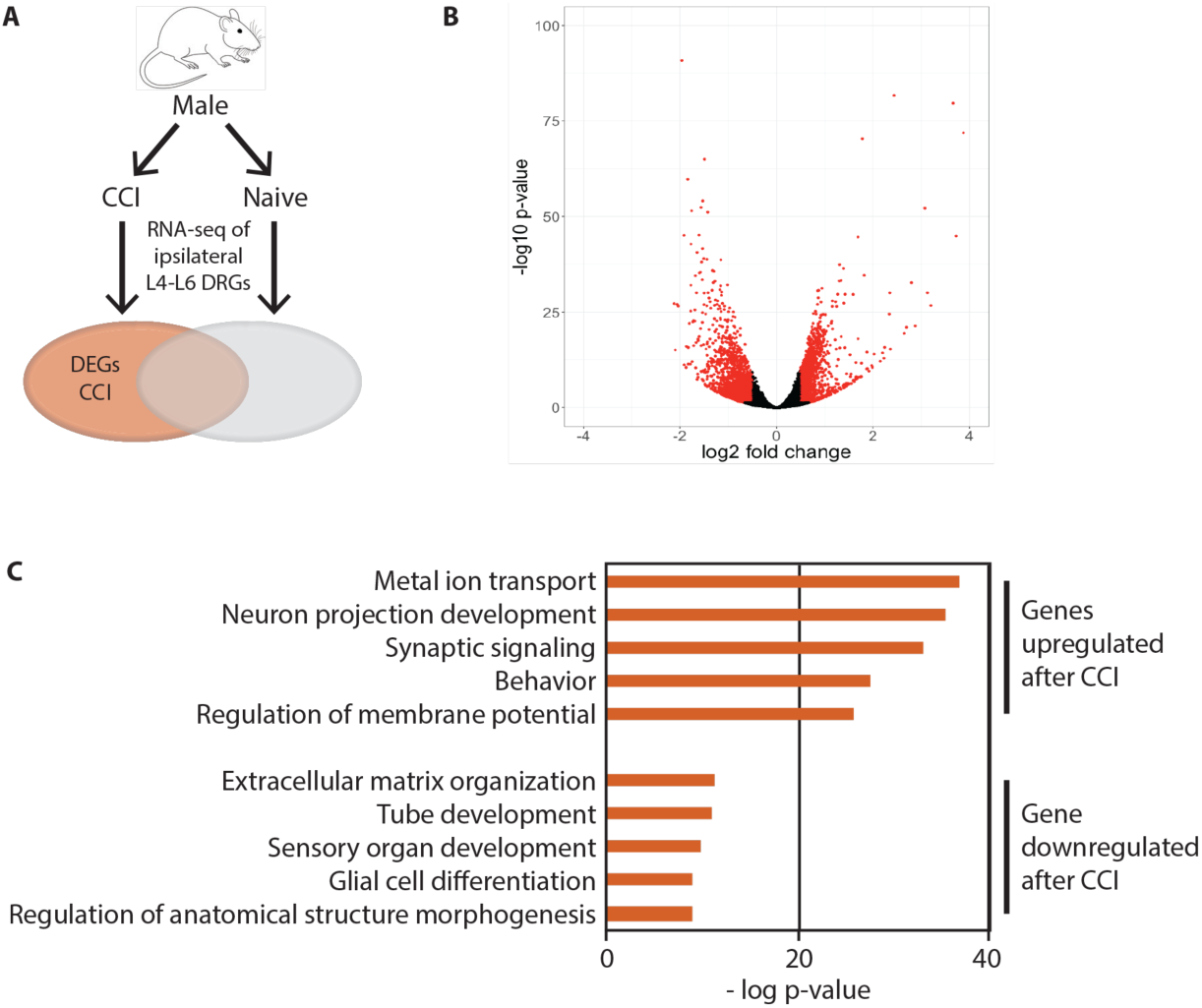
Differential gene expression between CCI and Naïve groups in male rats. A) Schematic diagram of experiment. Male rats were randomly assigned to the naïve group or receive CCI. RNA-seq performed on ipsilateral L4-L6 DRGs from each animal. Differentially expressed genes (DEG) are defined as genes expressed after CCI versus naïve with a |log_2_FC| > 0.5 and an adjusted p-value < 0.05. B) Volcano plot showing RNA-seq data of DRGs from naïve male rats and male rats following CCI. DEGs are designated in red. C) Gene ontology pathways associated with increased (top) and decreased (bottom) differential expression in CCI versus naive. DRG = dorsal root ganglia; CCI = chronic constriction injury; FC = fold change

**Table 1:** Top 25 differentially upregulated genes following CCI in males.

| Ensembl ID | Symbol | Gene description | log2 fold change | Standard error | Adjusted p-value |
| --- | --- | --- | --- | --- | --- |
| ENSRNOG00000004805 | *Stac2* | SH3 and cysteine rich domain 2 | 3.87 | 0.21 | 1.26E-72 |
| ENSRNOG00000018808 | *Vip* | Vasoactive intestinal peptide | 3.72 | 0.25 | 1.23E-45 |
| ENSRNOG00000006151 | *Reg3b* | regenerating islet-derived 3 beta | 3.66 | 0.19 | 2.14E-80 |
| ENSRNOG00000014327 | *Csrp3* | Cysteine and glycine rich protein 3 | 3.19 | 0.28 | 1.86E-27 |
| ENSRNOG00000010478 | *LOC299282* | Serine protease inhibitor | 3.12 | 0.26 | 9.23E-31 |
| ENSRNOG00000010079 | *Car3* | Carbonic anhydrase 3 | 3.07 | 0.19 | 6.46E-53 |
| ENSRNOG00000013496 | *Crisp3* | Cysteine-rich secretory protein 3 | 2.87 | 0.28 | 4.27E-22 |
| ENSRNOG00000056493 | *Mybpc1* | Myosin binding protein C, slow type | 2.79 | 0.22 | 2.03E-33 |
| ENSRNOG00000012703 | *Crh* | Corticotropin releasing hormone | 2.70 | 0.27 | 9.86E-22 |
| ENSRNOG00000018598 | *Ankrd1* | Ankyrin repeat domain 1 | 2.65 | 0.27 | 3.82E-20 |
| ENSRNOG00000003745 | *Atf* | activating transcription factor 3 | 2.44 | 0.12 | 2.22E-82 |
| ENSRNOG00000007017 | *Tmc2* | Transmembrane channel-like 2 | 2.36 | 0.28 | 5.33E-16 |
| ENSRNOG00000015156 | *Gal* | Galanin and GMAP prepropeptide | 2.35 | 0.20 | 9.47E-31 |
| ENSRNOG00000034258 | *Xirp2* | Xin actin-binding repeat containing 2 | 2.34 | 0.22 | 3.54E-25 |
| ENSRNOG00000000065 | *Pde6b* | Phosphodiesterase 6B | 2.24 | 0.26 | 1.83E-16 |
| ENSRNOG00000004583 | *Mb* | Myoglobin | 2.22 | 0.28 | 1.20E-13 |
| ENSRNOG00000020136 | *Tgm1* | Transglutaminase 1 | 2.20 | 0.26 | 5.05E-15 |
| ENSRNOG00000050004 | *Myot* | Myotilin | 2.15 | 0.28 | 7.75E-13 |
| ENSRNOG00000019447 | *Ecel1* | Endothelin converting enzyme-like 1 | 2.03 | 0.25 | 9.80E-15 |
| ENSRNOG00000015279 | *Vtcn1* | V-set domain containing T cell activation inhibitor 1 | 1.99 | 0.28 | 3.45E-11 |
| ENSRNOG00000029401 | *Actg2* | Actin, gamma 2, smooth muscle, enteric | 1.98 | 0.28 | 1.59E-11 |
| ENSRNOG00000046449 | *LOC100912228* | Pro-neuropeptide Y-like | 1.97 | 0.27 | 9.99E-12 |
| ENSRNOG00000010805 | *Fabp4* | Fatty acid binding protein 4 | 1.91 | 0.28 | 2.90E-10 |
| ENSRNOG00000048232 | *Plr747-ps* | Olfactory receptor pseudogene 1747 | 1.88 | 0.27 | 6.54E-11 |

**Table 2:** Top 25 differentially downregulated genes following CCI in males

| Ensembl ID | Symbol | Gene description | log2 fold change | Standard error | Adjusted p-value |
| --- | --- | --- | --- | --- | --- |
| ENSRNOG00000023824 | *RGD1561648* |  | -2.12 | 0.19 | 6.11E-28 |
| ENSRNOG00000025757 | *Myh6* | Myosin, heavy chain 6, cardiac muscle, alpha | -2.10 | 0.25 | 8.49E-16 |
| ENSRNOG00000017959 | *Cfap100* | Cilia and flagella associated protein 100 | -2.06 | 0.18 | 9.53E-28 |
| ENSRNOG00000014143 | *Col24a1* | Collagen, type XXIV, alpha 1 | -2.03 | 0.18 | 2.67E-27 |
| ENSRNOG00000016516 | *Mbp* | Myelin basic protein | -1.96 | 0.09 | 1.45E-91 |
| ENSRNOG00000059260 | *Aabr07053509.2* |  | -1.92 | 0.13 | 8.50E-46 |
| ENSRNOG00000021814 | *Tnfrsf25* | Tumor necrosis factor receptor superfamily, member 25 | -1.91 | 0.26 | 7.00E-12 |
| ENSRNOG00000048644 | *Ac115371.1* |  | -1.88 | 0.26 | 3.39E-11 |
| ENSRNOG00000033787 | *Adamts5* | ADAMTS-like 5 | -1.87 | 0.21 | 9.94E-17 |
| ENSRNOG00000012906 | *Bcas1* | Breast carcinoma amplified sequence 1 | -1.84 | 0.11 | 1.76E-60 |
| ENSRNOG00000016983 | *Myh7* | Myosin, heavy chain 7, cardiac muscle, beta | -1.83 | 0.21 | 1.95E-16 |
| ENSRNOG00000021242 | *Adam33* | ADAM metalloproteinase domain 33 | -1.83 | 0.25 | 4.15E-12 |
| ENSRNOG00000059605 | *Ddn* | Dendrin | -1.81 | 0.18 | 9.88E-23 |
| ENSRNOG00000018642 | *Leng8* | Leukocyte receptor cluster (LRC) member 8 | -1.78 | 0.24 | 4.13E-12 |
| ENSRNOG00000001325 | *Il3ra* | Interleukin 3 receptor subunit alpha | -1.78 | 0.16 | 5.46E-26 |
| ENSRNOG00000018603 | *Carns1* | Carnosine synthase 1 | -1.77 | 0.14 | 9.50E-33 |
| ENSRNOG000000062158 | *Rn60_1_2212.4* |  | -1.77 | 0.12 | 1.56E-43 |
| ENSRNOG00000017309 | *Rsrp1* | Arginine/serine-rich protein 1 | -1.75 | 0.11 | 3.09E-52 |
| ENSRNOG00000010438 | *Ct1b* | Carnitine palmitoyltransferase 1B | -1.74 | 0.23 | 9.55E-13 |
| ENSRNOG00000017508 | *Kmt5c* | Lysine methyltransferase 5C | -1.73 | 0.17 | 3.79E-23 |
| ENSRNOG00000007447 | *Pla2g4b* | phospholipase A2 group IVB | -1.71 | 0.16 | 1.72E-23 |
| ENSRNOG00000000563 | *Adamts14* | ADAM metalloproteinase with thrombospondin type 1 motif, 14 | -1.71 | 0.19 | 6.47E-17 |
| ENSRNOG00000007657 | *Col27a1* | Collagen, type XXVII, alpha 1 | -1.69 | 0.16 | 2.36E-23 |
| ENSRNOG00000013387 | *Tpcn2* | Two pore segment channel 2 | -1.69 | 0.15 | 2.71E-26 |
| ENSRNOG00000010626 | *Sphk1* | Sphingosine kinase 1 | -1.68 | 0.13 | 3.26E-35 |

### Differentially regulated genes in the dorsal root ganglion after nerve injury in female rats

To determine the effects of peripheral nerve injury on gene expression in female rats, we evaluated RNA-seq data at 14 days after CCI, and compared the expression profiles to those from the naive females (Figure 4A). In comparison to naive DRGs, 740 (3.6%) genes were differentially regulated after nerve injury (Figure 4B). Of these 740 differential genes, 480 genes (64.9%) were upregulated after injury and 260 genes (35.1%) were downregulated. The genes with the largest increase and decrease in gene expression following CCI are listed in Table 3 and Table 4, respectively. GO analysis identified that the top 5 enriched GO biological processes in the differentially expressed genes in females are shared between males and females following nerve injury (i.e., neuron projection development (GO:0031175), behavior (GO:0007610), regulation of membrane potential (GO:0042391)). However, even though males and females share enrichment in these biological processes, just 27-33% of these genes are common to both males and females (Supplemental figure 2). In addition to the three GO biological processes common to both males and females, the genes that were differentially expressed in females after injury were also enriched in 2 biological processes that were not present in males: Response to Cytokine (GO:0034097) and Response to Wounding pathways (GO:0009611) (Figure 4C). Among the 199 genes with increased expression in naive DRGs of female rats compared to the naïve male rats, 47 (23.6%) were downregulated and 15 (7.5%) were upregulated following CCI.

**Figure 4:**
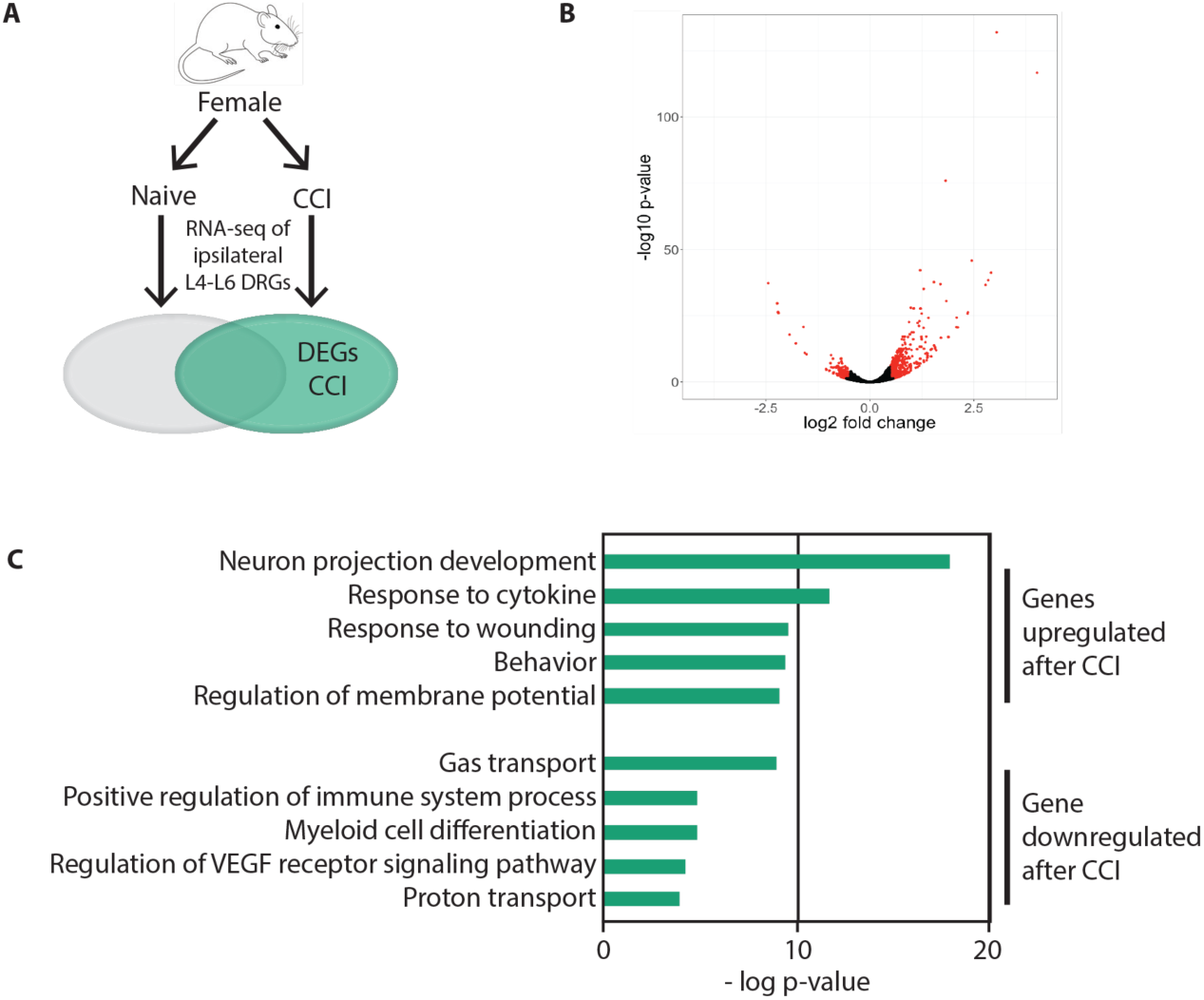
Differential gene expression between CCI and naïve groups in female rats. A) Schematic diagram of experiment. Female rats were randomly assigned to the naïve group or receive CCI. RNA-seq performed on ipsilateral L4-L6 DRGs from each animal. Differentially expressed genes (DEG) are defined as genes expressed after CCI versus naïve with a |log_2_FC| > 0.5 and an adjusted p-value < 0.05. B) Volcano plot showing RNA-seq data of DRGs from naïve female rats and male rats following CCI. DEGs are designated in red. C) Gene ontology pathways associated with increased (top) and decreased (bottom) differential expression in CCI versus naive. DRG = dorsal root ganglia; CCI = chronic constriction injury; FC = fold change

**Table 3:** Top 25 differentially upregulated genes following CCI in females

| Ensembl ID | Symbol | Gene description | log2 fold change | Standard error | Adjusted p-value |
| --- | --- | --- | --- | --- | --- |
| ENSRNOG00000004805 | *Stac2* | SH3 and cysteine rich domain 2 | 4.02 | 0.17 | 2.25E-117 |
| ENSRNOG00000003745 | *Atf3* | Activating transcription factor 3 | 3.05 | 0.12 | 1.21E-132 |
| ENSRNOG000000020136 | *Tgm1* | Transglutaminase 1 | 2.92 | 0.21 | 6.11E-42 |
| ENSRNOG00000001476 | *Cldn4* | Claudin 4 | 2.85 | 0.21 | 4.48E-39 |
| ENSRNOG000000013496 | *Crisp3* | Cysteine-rich secretory protein 3 | 2.78 | 0.21 | 2.51E-37 |
| ENSRNOG000000015156 | *Gal* | Galanin and GMAP prepropeptide | 2.45 | 0.16 | 1.78E-46 |
| ENSRNOG00000000065 | *Pde6b* | Phosphodiesterase 6B | 2.36 | 0.21 | 4.84E-27 |
| ENSRNOG000000018598 | *Ankrd1* | Ankyrin repeat domain 1 | 2.35 | 0.21 | 1.88E-26 |
| ENSRNOG000000058906 | *Ptprh* | Protein tyrosine phosphatase, receptor type, H | 2.12 | 0.21 | 2.64E-21 |
| ENSRNOG000000019447 | *Ecel1* | Endothelin converting enzyme-like 1 | 2.09 | 0.19 | 4.91E-25 |
| ENSRNOG000000015279 | *Vtcn1* | V-set domain containing T cell activation inhibitor 1 | 2.08 | 0.21 | 1.93E-21 |
| ENSRNOG000000006151 | *Reg3b* | Regenerating islet-derived 3 beta | 1.89 | 0.20 | 1.05E-17 |
| ENSRNOG000000001416 | *Vgf* | VGF nerve growth factor inducible | 1.84 | 0.15 | 3.03E-31 |
| ENSRNOG000000033531 | *Cacna2d1* | Calcium voltage-gated channel auxiliary subunit alpha2delta 1 | 1.82 | 0.10 | 1.15E-76 |
| ENSRNOG000000023892 | *Lce1f* | Late cornified envelope 1F | 1.71 | 0.19 | 2.21E-17 |
| ENSRNOG000000049882 | *Adcyap1* | Adenylate cyclase activating polypeptide 1 | 1.70 | 0.13 | 1.08E-37 |
| ENSRNOG000000050792 | *Tnfaip6* | TNF alpha induced protein 6 | 1.60 | 0.20 | 1.09E-12 |
| ENSRNOG000000005154 | *Fam150b* | Family with sequence similarity 150, member B | 1.58 | 0.20 | 2.74E-13 |
| ENSRNOG000000001338 | *Hpd* | 4-hydroxyphenylpyruvate dioxygenase | 1.57 | 0.19 | 8.99E-14 |
| ENSRNOG000000021029 | *Hamp* | Hepcidin antimicrobial peptide | 1.54 | 0.18 | 1.60E-14 |
| ENSRNOG000000004874 | *Flrt3* | Fibronectin leucine rich transmembrane protein 3 | 1.54 | 0.11 | 2.29E-38 |
| ENSRNOG000000046449 | *LOC100912228* | Pro-neuropeptide Y-like | 1.49 | 0.19 | 3.71E-13 |
| ENSRNOG000000017679 | *Cckbr* | Cholecystokinin B receptor | 1.48 | 0.21 | 2.68E-10 |

**Table 4:** Top 25 differentially downregulated genes following CCI in females

| Ensembl ID | Symbol | Gene description | log2 fold change | Standard error | adjusted p-value |
| --- | --- | --- | --- | --- | --- |
| ENSRNOG000000061299 | *LOC103694857* | Hemoglobin subunit beta-2 | -2.45 | 0.18 | 5.41E-38 |
| ENSRNOG000000045989 | *Hba-a3* | Hemoglobin alpha, adult chain 3 | -2.23 | 0.19 | 2.29E-30 |
| ENSRNOG000000047098 | *LOC103694855* | Hemoglobin subunit beta-2-like | -2.21 | 0.19 | 4.01E-27 |
| ENSRNOG000000058105 | *Hbb* | Hemoglobin subunit beta | -2.20 | 0.19 | 9.88E-27 |
| ENSRNOG000000047321 | *Hba-a2* | Hemoglobin alpha, adult chain 2 | -1.93 | 0.20 | 1.50E-18 |
| ENSRNOG000000029886 | *Hba-a1* | Hemoglobin alpha, adult chain 1 | -1.78 | 0.21 | 2.53E-15 |
| ENSRNOG000000034190 | *Ighm* | Immunoglobulin heavy constant mu | -1.60 | 0.16 | 1.62E-21 |
| ENSRNOG000000020951 | *Slc4a1* | Solute carrier family 4 (anion exchanger), member 1 | -1.57 | 0.21 | 9.82E-12 |
| ENSRNOG000000049829 |  | Aabr07060872.1 | -1.52 | 0.21 | 4.08E-11 |
| ENSRNOG000000000167 | *Alas2* | 5,-aminolevulinate synthase 2 | -1.06 | 0.21 | 1.38E-05 |
| ENSRNOG000000019183 | *Alox15* | Arachidonate 15-lipoxygenase | -1.05 | 0.21 | 2.25E-05 |
| ENSRNOG000000019660 | *Spib* | Spi-B transcription factor | -0.98 | 0.21 | 7.55E-05 |
| ENSRNOG000000024330 | *Ngp* | Neutrophilic granule protein | -0.97 | 0.20 | 5.48E-05 |
| ENSRNOG000000025757 | *Myh6* | Myosin, heavy chain 6, cardiac muscle, alpha | -0.95 | 0.15 | 7.90E-08 |
| ENSRNOG000000016516 | *Mbp* | Myelin basic protein | -0.93 | 0.13 | 7.21E-11 |
| ENSRNOG000000013228 | *Scrg1* | Stimulator of chondrogenesis 1 | -0.93 | 0.17 | 2.57E-06 |
| ENSRNOG000000017619 | *Aldh1a1* | Aldehyde dehydrogenase 1 family, member A1 | -0.89 | 0.13 | 1.66E-09 |
| ENSRNOG000000015085 | *Dmpk* | Dystrophia myotonica-protein kinase | -0.89 | 0.16 | 3.27E-06 |
| ENSRNOG000000060376 | *LOC100910237* | Uncharacterized LOC100910237 | -0.88 | 0.21 | 0.000658608 |
| ENSRNOG000000011917 | *Cd79b* | CD79b molecule | -0.87 | 0.19 | 0.000244955 |
| ENSRNOG000000010402 | *Hspb2* | Heat shock protein family B (small) member 2 | -0.85 | 0.16 | 2.83E-06 |
| ENSRNOG000000020296 | *LOC103693430* | Up-regulated during skeletal muscle growth protein 5 | -0.81 | 0.15 | 1.96E-06 |
| ENSRNOG000000010408 | *Polr2k* | Polymerase (RNA) II (DNA directed) polypeptide K | -0.80 | 0.15 | 3.44E-06 |
| ENSRNOG000000013593 | *Upk3a* | Uroplakin 3A | -0.80 | 0.19 | 0.000841337 |
| ENSRNOG000000032002 | *Hapln1* | Hyaluronan and proteoglycan link protein 1 | -0.78 | 0.14 | 9.02E-07 |
| ENSRNOG000000009768 | *Npy* | Neuropeptide Y | 1.47 | 0.19 | 7.89E-13 |
| ENSRNOG000000004033 | *Sema6a* | Semaphorin 6A | 1.41 | 0.13 | 6.32E-25 |

### Comparison of differentially regulated genes after nerve injury between males and females

To identify sex-specific transcriptional changes that occur following peripheral nerve injury, we compared the lists of differentially expressed genes after CCI between male and female rats (Figure 5A). We first visualized the extent of similarity/dissimilarity in the transcriptional profiles of the individual samples using the first two principal components from principal component analysis of all genes (Figure 5B). The first two principal components accounted for a total of 84% of the total variance among the samples and produced distinct clusters of the samples by treatment condition (i.e., naive, CCI) and sex. This segregation of samples was confirmed by hierarchical clustering of the top 200 genes that showed the greatest variability across all groups (Figure 5C). Of the 4056 genes that were differentially regulated between males and females, only 461 genes (11.4%) were common to both males and females regardless of direction of transcriptional changes (Figure 5D).

**Figure 5:**
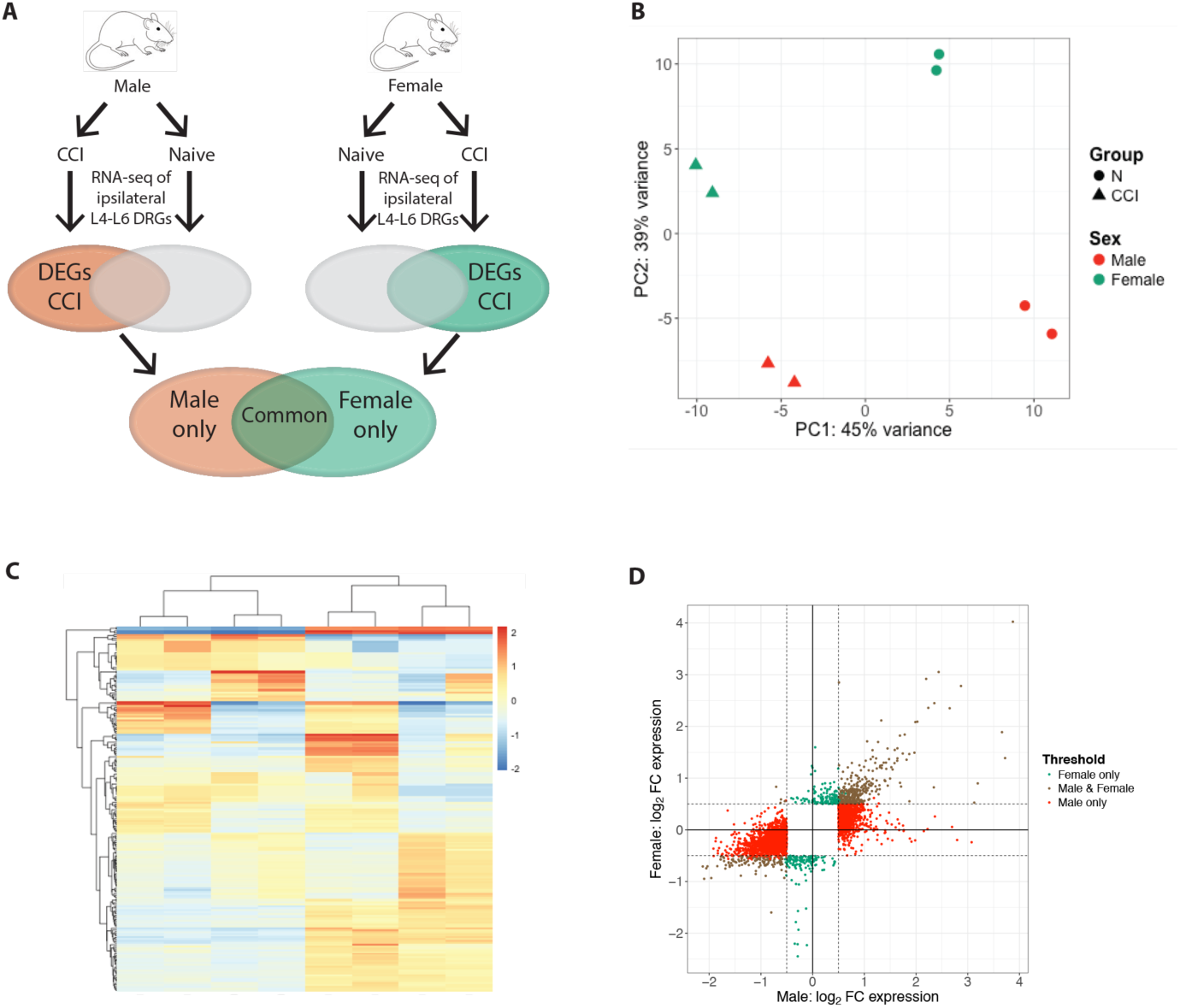
Comparison of differential gene expression between CCI and Naïve groups in male and female rats. A) Schematic diagram of experiment. Male and female rats were randomly assigned to the naïve group or receive CCI. RNA-seq performed on ipsilateral L4-L6 DRGs from each animal. Differentially expressed genes (DEG) defined as genes expressed after CCI versus naïve with a |log_2_FC| > 0.5 and an adjusted p-value < 0.05. B) Principal component analysis of male and female rats from the naïve and CCI groups. C) Heatmap that shows the log_2_FC of the 200 most differentially expressed genes between the CCI and Naïve groups in female and male rats. D) Log_2_FC expression between the CCI and naive males (x-axis) and females (y-axis). Threshold of |log_2_FC| > 0.5(dashed lines) with an adjusted p-value<0.05 designates DEGs in female rats only (green), male rats only (purple), and in both male and female rats (red). DRG = dorsal root ganglia; CCI = chronic constriction injury; FC = fold change

We then divided all upregulated genes that had an adjusted p-value <0.05 and |log_2_FC|>0.5 into the following 5 groups: 1) genes upregulated in both males and females, 2) genes upregulated in females with no change of gene expression in males, 3) genes upregulated in males with no change of gene expression in females, 4) genes upregulated in females and downregulated in males, and 5) genes upregulated in males and downregulated in females (Figure 6B). Of the 1429 genes that were upregulated in males, 336 (23.5%) were also upregulated in females. We used GeneMANIA to produce a co-expression network of these genes and identify functional pathways enriched in this gene set (Figure 6D). The top functional pathways identified in this set of upregulated genes were axon development, response to axon injury, regulation of neuron projection development, cellular calcium ion homeostasis, and sensory perception of pain. A total of 140 genes were specifically upregulated in females after CCI in that they had no significant change in gene expression in males. The functional pathways identified from this set of genes include response to glucocorticoid and response to corticosteriod (Figure 6D). Some examples of genes associated with these pathways include *Ahr*, *Cad*, *Cdkn1a*, *Fos*, *Reln*, and *Socs3* (Figure 6F).

**Figure 6:**
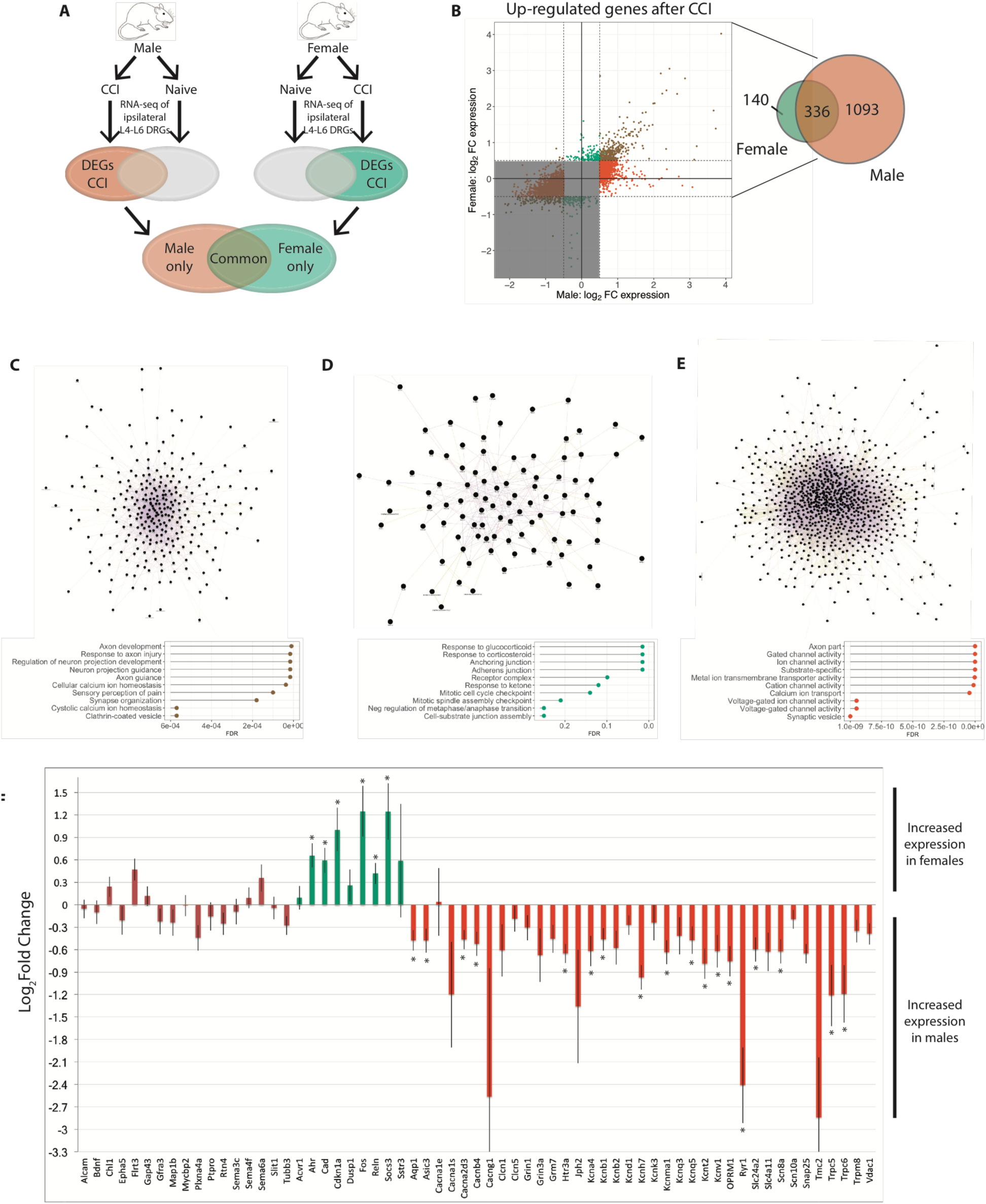
Co-expression networks of differentially expressed genes. A) Schematic diagram of experiment. Male and female rats were randomly assigned to the naïve group or receive CCI. RNA-seq performed on ipsilateral L4-L6 DRGs from each animal. Differentially expressed genes (DEG) defined as genes expressed after CCI versus naïve with a |log_2_FC| > 0.5 and an adjusted p-value < 0.05. B) Log_2_FC expression between the CCI and naive males (x-axis) and females (y-axis) for DEGs upregulated in CCI versus naive. Threshold of |log_2_FC| > 0.5 (dashed lines) with an adjusted p-value<0.05 designates DEGs in female rats only (green), male rats only (red), and in both male and female rats (brown). Venn diagram shows the numbers of DEGs identified in each of these groups. C-E) Co-expression network for differentially expressed genes in both male and female rats C), female rats only D), and male rats only E) with increased gene expression (top) obtained from GeneMANIA. Functional pathway analysis lists the top 10 gene ontology pathways (middle) with the FDR for each term. F) For each of the top-ranking pathways, the log_2_FC expression values for DEGs common to several of these pathways are plotted (bottom). Genes that significantly differ between males and females with an adjusted p-value<0.05 are indicated with an asterisk. Positive values of the log_2_FC indicate increased expression in females compared to males and negative values indicate increased expression in males versus females. DRG = dorsal root ganglia; CCI = chronic constriction injury; FDR = false discovery rate; FC = fold change

A total of 1093 genes were specifically upregulated in males after CCI which had no expression change in females. The functional pathways identified from this set of genes include gated channel activity (GC:002836), ion cannel activity (GO:0022839), substrate-specific channel activity (GO:0022838), and metal ion transmembrane transporter activity (GO:0046873) (Figure 6E). Examples of genes associated with these pathways include potassium channels (i.e., *Kcna4*, *Kcnb1*, *Kcnh7*, *Kcnma1*, *Kcnq5*, *Kcnt2*, *Kcnv1*), calcium channels (i.e., *Cacna2d3*, *Cacnb4*), *Oprm1*, and *Scn8a* (Figure 6F). A complete list of genes found to be significantly differentially expressed between males and females after CCI is provided in Supplemental Table 4. Four genes were regulated in opposite directions after injury. *Adamts4*, *Cyp2s1*, *Top2a*, and *Anln* were upregulated in female rats and downregulated in male rats after CCI compared with naive controls. Of note, no genes were identified that were upregulated in males and downregulated in females.

We divided the genes downregulated after CCI that had an adjusted p-value <0.05 and | log_2_FC |>0.5 into the following 3 groups: 1) genes downregulated in both males and females, 2) genes downregulated in females with no change of gene expression in males, 3) genes downregulated in males with no change of gene expression in females (Supplemental figure 3B). No significant functional pathways were enriched using the genes common to both males and females. Genes downregulated only in females showed significant enrichment in the Mitochondrial membrane part GO cellular component (Supplemental figure 3C). Genes downregulated only in males showed significant enrichment in pathways enriched for extracellular matrix and the Wnt signaling pathway (Supplemental figure 3D).

Supplemental figure 4 shows the results of biologic validation of 10 genes by qPCR using a separate cohort of animals. We first identified an appropriate endogenous control gene by evaluating 6 candidate reference genes that showed the most stable gene expression among naïve and injured DRG of both sexes. The NormFinder algorithm calculated the lowest stability score for *Hmbs* which is consistent with stable expression among conditions (Table 5). Due to missing data from one sample *Ywhaz* could not be included in this analysis. The relative changes in gene expression derived by qPCR were in agreement with those detected by RNA-seq.

**Table 5:**
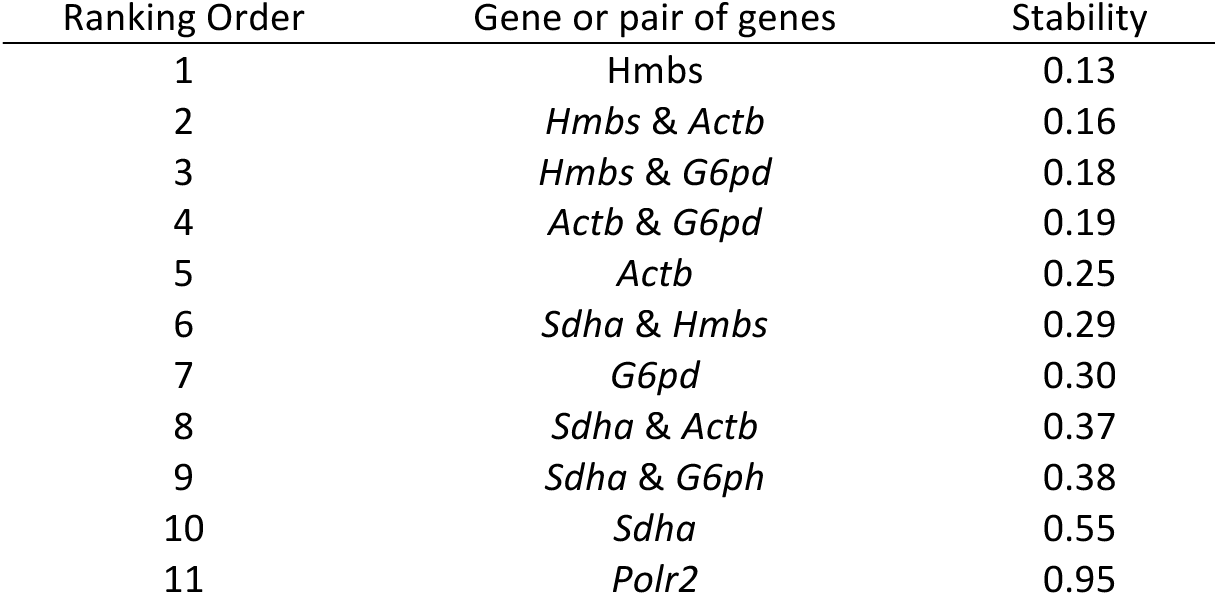
Stability values of candidate reference genes in the DRG as determined by NormFinder

## DISCUSSION

Few studies have addressed sex-specific alterations in gene expression in neuropathic pain models, and as a result, the molecular pathways underlying sex differences in neuropathic pain are poorly understood. In addition, gene expression data are available for several types of neuronal tissue from rats, mice, and humans; however, to our knowledge, RNA-seq derived whole transcriptome analysis of the PNS has not been performed in these organisms. In the present study, we use RNA-seq to compare the gene expression profiles in lumbar DRGs between male and female rats under both naïve and nerve injury conditions. Our results highlight vast differences in the regulation of genes in PNS that are associated with pain-relevant pathways between females and males after injury. However, both sexes upregulate similar groups of genes associated with axon regeneration and neuron regrowth pathways.

### Sex differences in gene expression in the naïve DRG

Our gene expression profiles from naive DRGs of male and female rats provide important insights into the normal expression of PNS genes in an unperturbed organism. Remarkably, only 4.6% of all genes expressed in the naïve DRGs were differentially expressed by sex. Furthermore, these genes that were differentially expressed between sexes did not enrich for pathways directly implicated in either nerve injury or nociception. However, genes that had increased expression in naïve females did show enrichment in immune-related pathways. Increased expression in females of immune-related genes is not unexpected since the connection between sex hormones and immune function is tightly linked. While comprehensive mechanisms for the sexual dimorphism in nociceptive pain have yet to be determined, current evidence supports an important role of sex hormones in pain modulation through the innate and adaptive immune systems (Rosen et al. 2017; Sorge and Totsch 2017). Overall, females mount a stronger humoral, cellular, and inflammatory response than males which is largely attributed to differences in circulating levels of sex hormones especially the decreased levels of androgens (Gaillard and Spinedi 1998; Bouman et al. 2005). The effects of circulating androgens are known to dampen proinflammatory cytokine production through release of IL-10 by Th1 cells (Bebo et al. 1999) in males. Inflammatory cytokines modulate neuronal excitability through changes in the composition, density, and spatial distribution of ion channels and receptors in the neuronal membrane (Inoue et al. 1999; von Banchet et al. 2005). Therefore, males may experience less pain as a result of a damped pro-inflammatory immune response after a pain-initiating stimulus. Indeed, evidence from experimental pain studies of pain-free individuals demonstrates a more pronounced temporal summation and greater pain sensitivity in females versus males (Fillingim et al. 2009).

### Similar gene expression changes after peripheral nerve injury between males and females

Genes that were similarly upregulated in the DRG of both sexes after nerve injury were related to neuron regeneration, pain, and intercellular signaling. Following nerve injury, neuronal pathways involved in the regeneration of damaged axons are activated in attempts to recover lost motor, sensory, and autonomic functions. These regenerative mechanisms involve sprouting from the end of the damaged axon toward the denervated segments of the cell, collateral branching of the undamaged neurons around the injury, and long-term reorganization of neuronal circuits in the CNS in response to aberrant peripheral signals (Hanz and Fainzilber 2006). Changes in gene expression and redistribution of receptors and ion channels in the neuronal membrane (Scholz and Woolf 2002) may partially compensate for incomplete reinnervation; however, altered sensation and loss of fine motor control may produce functional deficits as well as maladaptive alterations (e.g., enhanced spinal reflexes, neuropathic pain).

The few existing studies of sex differences in nerve regeneration suggest that regeneration of myelinated neurons occurs more rapidly in females than males (Kovacic et al. 2003; Kovacic et al. 2004; Kovacic et al. 2009). However, Tong and colleagues recently found no sex differences in axon regeneration of axotomized neurons (Tong et al. 2015). Our findings support the latter study since after CCI no significant sex-specific transcriptional differences exist in genes known to be involved in axon regeneration and neuron projection development. These discrepancies may be attributed to different nerve injury models, measures of regeneration, and timing of assessments.

### Gene expression profiles after injury differ between males and females

After nerve injury, male and female rats showed transcriptional changes in genes that share common GO biological processes (i.e., Neuron Projection Development, Behavior, and Regulation of Membrane Potential). However, less than one third of the genes that were enriched for each of these common processes are shared between males and females. For example, *Akap9* (A-kinase anchor protein 9) is a member of the Regulation of Membrane Potential GO biologic process and, in our study, is significantly upregulated in females but not in males after CCI. *Akap9* interacts with members in signaling transduction pathways (e.g., protein kinase A, serine/threonine kinase protein kinase N, protein phosphatase 1) (Takahashi et al. 1999) and loss-of-function mutations are associated with Long QT syndromes (Chen et al. 2007). The role of *Akap9* in peripheral nerves has not been reported; however, its link to regulating membrane potential suggests *Akap9* could contribute to a sex-specific pain mechanism by increasing hyperexcitability after nerve injury in females.

*Oprm1* is also represented in the Regulation of Membrane Potential GO biologic process, but in contrast to *Akap9*, *Oprm1* is significantly upregulated after injury in males but not in females. Importantly, *Oprm1* encodes the μ-opioid receptor (MOR) which is targeted by the agonist morphine to produce profound analgesia. Zubieta and colleagues (Zubieta et al. 2002) found that, in relation to women, men experienced increased activation of the endogenous analgesic mechanisms in several brain regions during infusion of hypertonic saline into the masseter muscle as demonstrated by increased binding of radiolabeled carfentanil to MORs. Further, several preclinical studies report that morphine produces greater analgesia in male animals in the setting of chronic inflammatory pain (Loyd and Murphy 2006; Wang et al. 2006). In the present DRG-focused study, male rats had significantly higher *Oprm1* expression than females after injury, which suggests a previously unidentified peripheral mechanism that may also contribute to sex differences in morphine analgesia and endogenous analgesic mechanisms. Consistent with this finding, prior work in our lab has demonstrated a relative decrease in efficacy of the peripheral-acting MOR agonist DALDA (dermorphin [D-Arg2, Lys4 (1-4) amide) in neuropathic pain-related behavior after systemic administration in female rats (Tiwari et al. 2016).

Another important DRG transcript we found upregulated after CCI in both male and female rats was *Csf1* (colony-stimulating factor 1). *Csf1* is transported from the DRG to the spinal cord dorsal horn where it binds to its receptor (i.e., *Csf1r*) located on microglia whose activation promotes the CNS changes necessary for mechanical hypersensitivity (Guan et al. 2016). Our results are consistent with a recent study (Guan et al. 2016) in which *Csf1* was reported to be expressed *de novo* in injured sensory neurons following peripheral nerve injury. However, we found *Csf1* expression was 1.7 times higher in females than in males. Further studies will be required to examine if and how sex-specific regulation of *Csf1* factors into differences between males and females in their susceptibility to chronic pain syndromes.

A limitation of our findings is our inability to resolve cell type specificity of gene expression from bulk DRG. Recent single cell expression studies have identified subpopulations of neuronal and non-neuronal cells within male DRG (Usoskin et al. 2015; Li et al. 2016). Ultimately, understanding how sex-specific differences in PNS gene expression arise at a single cell level should help facilitate drugs regimens that limit off-target effects.

Our findings build upon existing knowledge of differential pain sensitivity and susceptibility to chronic pain and highlight the profound differences in peripheral pain mechanisms between males and females after nerve injury. First, baseline differences in immune function in the PNS may predispose females to developing chronic pain conditions. Second, functional pathways relevant to both sexes that become altered after injury may have important underlying differences in gene expression. Failures of preclinical treatments to translate into humans may be at least partially due to sex-specific alterations in gene expression and existing biases for exclusion of females in preclinical studies. Increased attention to sex differences in preclinical pain studies may improve the translational relevance to clinical populations.

## METHODS

### Animal models

Adult male and female Sprague Dawley rats (Harlan Bioproducts for Science, Indianapolis, IN) were randomly assigned to undergo CCI surgery or no treatment (i.e., naive control). All surgical procedures were performed by the same individual to avoid variation in technique. *All animals*: Animals were allowed to acclimate for a minimum of 48 hours before any experimental procedures, housed 2-3 per cage, and given access ad libitum to food and water. *CCI surgery group*: CCI surgery to the sciatic nerve was performed as previously described (Bennett and Xie 1988). Under 2% isoflurane, a small incision was made at the level of the mid-thigh. The sciatic nerve was exposed by blunt dissection through the biceps femoris. The nerve trunk proximal to the distal branching point was loosely ligated with four 4-0 silk sutures placed approximately 0.5 mm apart until the epineuria was slightly compressed and minor twitching of the relevant muscles was observed. The muscle layer was closed with 4-0 silk suture and the wound closed with metal clips. On postoperative day 14, naive and CCI rats were euthanized by overdose of isoflurane and the ipsilateral L4-L6 DRGs were quickly dissected, immediately submerged in liquid nitrogen, and stored at -80°C until RNA extraction. The ipsilateral L4-L6 DRGs from a single rat were pooled and define each sample. Of note, we did not identify the stage of estrus in the female rats. Existing literature suggests that freely cycling rodents do not exhibit increased variability of study outcomes including pain behaviors (Mogil and Chanda 2005; Prendergast et al. 2014; Klein et al. 2015). All procedures involving animals were reviewed and approved by the Johns Hopkins Animal Care and use Committee and were performed in accordance with NIH Guide for the Care and Use of Laboratory Animals.

### Behavior testing

Mechanical hypersensitivity was measured using von Frey monofilaments as previously described (Chaplan et al. 1994). Animals were assessed at baseline and on day 14 after CCI (Supplemental figure 1). The animals were placed in individual plexiglass cages on top of a wire mesh and allowed to acclimate for 1 hour. Tactile stimulation to the midplantar surface of each hind paw was performed using calibrated monofilaments with gradually increasing stiffness (i.e., 0.37, 0.61, 1.23, 2.0, 4.0, 5.93, 10.0, 13.5 grams). Each monofilament was applied for 4-6 seconds in the area between the footpads on the plantar surface of the hind paw. Monofilaments with increasing force were applied until a positive response was observed (e.g., abrupt paw withdrawal, shaking, licking). When a positive response was observed, the monofilament with the next lower force was then applied. The test continued 1) for 5 stimulations after a positive test was observed or 2) the upper or lower range of the von Frey monofilament set was obtained. The paw withdrawal threshold was determined as previously described. If a rat did not achieve at least a 50% reduction in baseline paw withdrawal threshold after 48 hours or on day 14, then this animal was not used (n=3). In addition to one animal of each sex that failed to show adequate reduction of withdrawal thresholds, one naïve animal was remove from analysis due to responses at a much lower mechanical threshold than normal.

### RNA isolation

Total RNA was extracted from pooled ipsilateral lumbar DRGs (L4-6) from one rat using the Qiagen RNeasy Mini Prep Kit (Qiagen, Valencia, CA; #74104) with on-column DNase digestion (Qiagen; #79254) according to manufacturer’s instructions. RNA concentration was measured using the Nanodrop ND-2000 Spectrophotometer (Thermo Fisher Scientific, Waltham, MA) and RNA integrity was assessed using RNA Nano Eukaryote chips in an Agilent 2100 Bioanalyzer (Agilent Technologies, Palo Alto, CA).

### Library construction and sequencing

Five hundred nanograms of total RNA per sample was used to construct sequencing libraries (n=1 rat/sample and run in independent biological duplicates per group per sex). Strand-specific RNA libraries were prepared using the NEBNext Ultra Directional RNA Library Prep Kit for Illumina (New England Biolabs; # E7420S) after poly(A) selection by the NEBNext poly(A) mRNA Isolation Module (New England Biolabs; #X7490) according to manufacturer’s instructions. Samples were barcoded using the recommended NEBNext Multiplex Oligos (New England Biolabs; #E7490). Size range and quality of libraries were verified on the Agilent 2100 Bioanalyzer (Agilent Technologies; Palo Alto, CA). RNA-seq libraries were quantified by qPCR using the KAPA library quantification kit (KAPA Biosystems). Each library was normalized to 2nM and pooled in equimolar concentrations. Paired-end sequencing was performed on an Illumina HiSeq2500 (Illumina, San Diego, CA). Two independent biological replicates of 1 rat per group per sex were run for a total of 8 libraries. Libraries were sequenced to an average depth of 38 million reads per sample.

### Data analysis

Sequencing reads were aligned to the rat reference genome (rn6) using HISAT2 (Kim et al. 2015). Aligned reads were filtered to remove ribosomal RNA and visualized using the Integrative Genomics Viewer (Robinson et al. 2011). A gene count matrix that contained raw transcript counts for each annotated gene were generated using the *summarizedOverlaps* function of the GenomicAlignments package in R (Lawrence et al. 2013) against the Ensemble rn6 transcriptome. We filtered this matrix for low count genes by retaining only those genes with >5 reads in all 4 samples for each sex.

To identify genes that were differentially regulated following nerve injury, transcript counts were normalized and log_2_ transformed using the default normalization procedures in DESeq2 (Love et al. 2014). The differential expression analysis was first performed separately for each sex using default parameters. This analysis identified differentially expressed genes between the naive and CCI groups within males or females. The interaction of sex on differential gene expression after injury was evaluated by the interaction term included in the design matrix within DESeq2. All downstream analyses on RNA-seq data were performed on data obtained from DESeq2. An adjusted p-value <0.05 and an absolute log_2_ fold change > 0.5 were used to define differentially expressed transcripts between naive and injured animals. Genes with differential expression between groups were then included in pathway analysis to infer their functional roles and relationships.

Gene ontology analysis for enriched biological processes in each set of differentially enriched genes identified by DESeq2 was performed using Metascape [http://metascape.org] (Tripathi et al. 2015) with a minimum enrichment of 1.0 and a p-value cutoff of 0.05. Metascape compiles data monthly from publicly available resources (e.g. NCBI, Reactome, GO, KEGG) to provide comprehensive analysis of a list of genes. All significant differentially enriched genes identified by DESeq2 were used to construct network-based functional associations using the GeneMANIA algorithm (Warde-Farley et al. 2010) as a plug-in within Cytoscape version 3.4.0 (Montojo et al. 2010).

### Validation of RNA-seq by qPCR

Total RNA extracted from DRGs of a separate cohort of rats was used to confirm the relationship of gene expression trends between sexes in selected genes by qPCR. As previously described, first-strand cDNA synthesis from 500ng total RNA in a 20μl reaction was performed using random hexamer primers and the SuperScript III Reverse Transcriptase (ThermoFisher Scientific) according to manufacturer’s instructions. cDNA was diluted 1:4 with nuclease-free water and stored at -20°C. mRNA sequence for each gene was retrieved from NCBI. Forward and reverse primers for each gene were designed using the PrimerQuest Tool (IDT, Coralville, Iowa) to span one or more introns. Primers were obtained through Integrated DNA Technologies (IDT, Coralville, Iowa) and sequences are provided in Supplemental Table 1.

Each 20 μl qPCR reaction consisted of 10ul 2X Power SYBR Green Master Mix (Thermo Fisher Scientific, Waltham, MA), 200 nM each forward and reverse primer, and 2 μl diluted cDNA. PCR of each target was performed using the 7900HT Fast Real-Time PCR system (Applied Biosystems, Waltham, MA) with the following thermocycling conditions: initial denaturation at 95°C for 10 minutes followed by 40 cycles of 95°C for 10 seconds and 60°C for 60 seconds. Two biological replicates were assayed for each group and each biological replicate was run in triplicate for each target gene. Nuclease-free water was included in each plate as a no-template control.

PCR efficiencies of each primer set were determined using the slope of standard curves constructed with Cq values obtained from 5-fold serial dilutions of pooled cDNA from DRGs of each group (e.g., male naïve, female naïve, male CCI, female CCI). The efficiency was calculated using they formula: E=10^−1/slope^. Dissociation curve analysis was used to identify amplification of non-specific products including primer dimers.

### Identification of an endogenous control

To identify a stable endogenous control gene for normalization of the target genes in qPCR we selected six candidate reference genes (i.e., *Sdha*, *Hmbs*, *Polr2a*, *G6pd*, *Ywhaz*, *Actb*) that showed little variation in the RNA-seq data. The expression stability of each candidate reference gene across groups was analyzed using the NormFinder function in R (https://www.moma.dk/normfinder-software) (Andersen et al. 2004). For each candidate gene, NormFinder calculates a stability value for each gene based on the genes expression variation among samples within the same group and variation between different group. This stability value enables candidate genes to be ranked according to their expression stability among different experimental conditions. The gene or pair of genes with the lowest stability value (i.e., highest expression stability) was selected as the endogenous control for relative gene expression calculations of each target gene.

## Statistical analysis

### Differential gene expression analysis

Several pairwise comparisons were performed. First, we compared gene expression in the DRGs from naive rats in male and females to identify transcripts that are expressed at significantly higher or lower levels than in the opposite sex. Then, we identified transcripts that were differentially regulated following injury compared with the naive tissue within the same sex (i.e., male CCI versus male naive tissue, female CCI versus female naïve tissue). Last, we compared genes that were differentially regulated following injury between males and females.

### Quantitative real-time PCR

Default settings were used to define quantification cycle (C_q_) values using SDS software version 2.4.1 (Applied Biosystems, Waltham, MA). The Cq values were averaged over three technical replicates. If the standard deviation of this average was > 0.20, the outlying replicate was removed and the Cq was averaged over the two remaining technical replicates. The 2^−ΔΔCT^ method was used to convert C_q_ values into relative gene expression for each gene (Livak and Schmittgen 2001).

## ACKNOWLEDGEMENTS

We thank members of the Taverna and Guan labs for critical reading of this manuscript and Rakel Tryggvadóttir and Colin Callahan for their technical assistance. This study was supported by the Blaustein Pain Research and Education Endowment (K.E.S., S.D.T.) and National Institutes of Health grants R01GM106024 (S.D.T.), R01GM118760 (S.D.T.), F32NR015728 (K.E.S.), R01NS070814 (Y.G.), and R21NS099879 (Y.G.), and R01HG006282 (H.J.).

## AUTHOR CONTRIBUTIONS

KES, Conception and design, Acquisition of data, Analysis and interpretation of data, Drafting or revising the article; WZ, Analysis and interpretation of data, Drafting or revising the article; ZJ, Analysis and interpretation of data; SH, Acquisition of data; HJ, Analysis and interpretation of data, Drafting or revising the article; YG, Conception and design, Analysis and interpretation of data, Drafting or revising the article; SDT, Conception and design, Analysis and interpretation of data, Drafting or revising the article.

